# TadA-Derived Cytosine Base Editor for Precise Genome Editing in Zebrafish

**DOI:** 10.1101/2025.01.28.635338

**Authors:** Wei Qin, Sheng-Jia Lin, Cassidy Petree, Pratishtha Varshney, Gaurav K. Varshney

**Affiliations:** Genes & Human Disease Research Program, Oklahoma Medical Research Foundation, Oklahoma City, OK, USA

## Abstract

CRISPR base editors are crucial for precise genome manipulation. Existing APOBEC-based cytosine base editors (CBEs), while powerful, exhibit indels and sequence context limitations, where TC-context preferences restrict effective editing of CC and GC motifs. To address these challenges, we evaluated various tRNA adenine deaminase (TadA)-derived CBEs, ultimately engineering zTadCBE that demonstrates high editing efficiency, minimized off-target effects, and an expanded targeting range. Our approach integrates beneficial mutations from TadA-based adenine base editors (ABEs) with SpRYCas9n-enhanced protospacer-adjacent motif (PAM) compatibility. Additionally, we engineered expanded-window zTadCBE variants, zTadCBE-ex1 and zTadCBE-ex2, to target wider nucleotide ranges, further increasing the versatility of this tool. To demonstrate the utility of zTadCBE variants in the functional assessment of genetic mutants, we generated a model for CDH23-associated hearing loss to validate the pathogenicity of a patient-specific variant in zebrafish. Furthermore, we induced a premature stop codon in the mediator complex gene *med12* to inactivate its function using a CRISPR-STOP strategy and recapitulated patient-specific phenotypes in the founding (F0) generation. zTadCBE variants thus offer a robust set of CBEs for precise and efficient C-to-T editing in zebrafish, promising to advance the rapid functional assessment of genetic variants in vivo.

## Introduction

Genome editing has experienced a remarkable breakthrough with the development of clustered regularly interspaced short palindromic repeat (CRISPR) base editors, a revolutionary technology in molecular biology that enables the precise and efficient modification of single nucleotides within the genome, without introducing double-stranded breaks. This technology offers powerful capabilities for studying genetic variants in cell lines or animal models, effectively mimicking certain human genetic diseases or variants of unknown significance (VUS). Base editors comprise a catalytically inactive CRISPR-associated protein 9 (Cas9) fused to a deaminase enzyme and fall into two primary classes: cytosine base editors (CBEs) and adenine base editors (ABEs). CBEs use a cytidine deaminase such as rat APOBEC1 (rAPOBEC-1) that converts cytosine (C) to uracil (U)—which is read as thymine (T) during DNA replication or repair—and a uracil glycosylase inhibitor (UGI) that prevents C-to-U reversions by the cell’s natural repair mechanisms^1^. On the other hand, ABEs use a laboratory-evolved tRNA adenine deaminase (TadA) that specializes in converting adenine (A) to guanine (G), or its complementary base thymine (T) to cytosine (C)^2^.

Several CBEs, such as BE4, BE4-Gam, BE4max, evoFERNY-BE4max, and evoAPOBEC1-BEmax, have been tested in cells, plants, and various animal models^2–7^. While they offer great promise for genome editing, they show variable or limited editing efficiencies, substantial off-target editing, and induce significant insertions and deletions (indels) in zebrafish and many other model organisms^8–10^. Even the most advanced and efficient CBEs such as BE3, BE4, BE4max, and AncBE4max—all of which harbor an rAPOBEC1 deaminase domain^1,2,7,8,11,12^—suffer from sequence context preferences that limit the genomic sites they can target; APOBEC1 has a strong preference to edit the C in T<u>C</u> motifs rather than in G<u>C</u> motifs, and these biases are more pronounced in zebrafish compared to human cell lines^8,13^.

Many groups have optimized CBEs using different cytosine deaminases and engineered variants (e.g., hAPOBEC3A, PmCDA1, hAID, and Anc689) to reduce off-target DNA edits and broaden editing scope^7,14–16^. For example, the improved editing efficiency of AncBE4max comes from the use of modified nuclear localization sequences (NLSs) to enhance its nuclear entry, as well as a codon-optimized version of the ancestral deaminase Anc689 APOBEC^7^. EvoFERNY-BE4max is a fourth-generation CBE using a directed evolution-optimized APOBEC1 deaminase (evoFERNY) fused to a Cas9 nickase^6^, imparting superior editing capabilities. Despite these optimizations, challenges to achieving efficient and precise edits remain.

In contrast to cytidine deaminases, TadA enzymes used by ABEs are relatively less processive and induce A-to-G DNA conversions with over 99.9% purity, minimal indels, and a compact editing window, allowing current ABE variants, like ABE8e (also called TadA-8e), to achieve higher editing efficiencies, greater single-nucleotide precision, and lower Cas-independent off-target editing than CBEs^17^. New research has therefore focused on re-engineering the TadA enzyme from TadA-8e to produce artificial variants that deaminate cytosine instead of adenine, overcoming the challenges posed by CBEs^18–20^. Based on TadA engineering, multiple variants of Tad-CBEs, such as TadCBEa, TadCBEd, TadCBEd_V106W, TadDE, eTd-CBE, Td-CBEmax, and CBE-T1.14, have been generated that can mediate efficient C-to-T conversions in human cells and plants^18–22^. Recently, we developed a next-generation TadA-based ABE variant, ABE-ultramax, with up to 100% A-to-G editing efficiency^23^. However, the application of TadA variants in zebrafish—a key model system for studying human genetic diseases, owing to its genetic versatility, rapid development, and similarities to mammals— remains limited. Successful demonstrations of CBEs are restricted to a few loci, with only zAncBE4-max and BE4-Gam tested with variable efficiencies^3,8,11,24^, leaving the potential of these variants for optimizing zebrafish CBEs largely unexplored.

Here, we comprehensively evaluated existing CBEs, including TadA-derived variants, for cytosine to thymine editing in zebrafish. Our analysis revealed that TadCBEa mediates highly efficient C-to-T editing, while TadCBEmax induces the fewest indels. Building on these findings, we developed a novel TadA-derived CBE, zTadCBE, which combines high editing efficiency with low indel rates. Our application of this tool to develop zebrafish disease models further underscored its practical utility, enabling the targeting of previously inaccessible loci. These results highlight the significant potential of zTadCBE for expanding the scope of functional genomics and disease modeling in zebrafish.

## Results

### Evaluation and Comparison of the in vivo Editing Efficiencies of Six Tad-A-derived CBEs in Zebrafish

To assess the efficacy of TadA-derived cytosine base editors (CBEs) in converting cytosine (C)-to-thymine (T) in zebrafish, and to systematically compare their editing efficiency with existing CBEs, we selected six representative CBE systems from diverse research groups, three TadA-derived systems—TadCBEmax, TadCBEa, CBE-T1.14, and three existing systems—AncBE4max, evoAPOBEC1, evoFERNY. To comprehensively capture their editing characteristics, we selected six endogenous loci with distinct sequence context preferences. To reduce variability arising from differences in codon usage, we constructed all CBE tools using zebrafish codon optimization (**Fig. 1a**), with each construct containing N-terminal and C-terminal bipartite nuclear localization signals (bpNLS). To analyze base editing outcomes, we injected 2’-O-methyl-3’-phosphorothioate (MS)-modified single guide RNAs (sgRNAs) into one-cell stage zebrafish embryos, extracted genomic DNA at 48 h post-fertilization (hpf), and performed next-generation sequencing (NGS) following polymerase chain reaction (PCR) amplification of target regions. Our NGS data revealed that all six CBE systems achieved effective C-to-T base editing in zebrafish, albeit with varying efficiency (**Fig. 1b**). Across all six target sites of all CBEs evaluated, TadCBEa exhibited the highest efficiency, followed by TadCBEmax, while AncBE4max had the lowest efficiency (**Fig. 1c**). Additionally, analysis of insertions and deletions (indels) indicated that TadCBEmax sustained high on-target activity with relatively low indel levels, reflecting an optimal balance between high editing efficiency and minimal off-target effects (**Fig. 1d and 1e**). Collectively, we demonstrated that each TadA-derived CBE variant has certain limitations, necessitating their optimization to achieve specific C-to-T editing with low indel levels and high-on target activity.

**Fig. 1:**
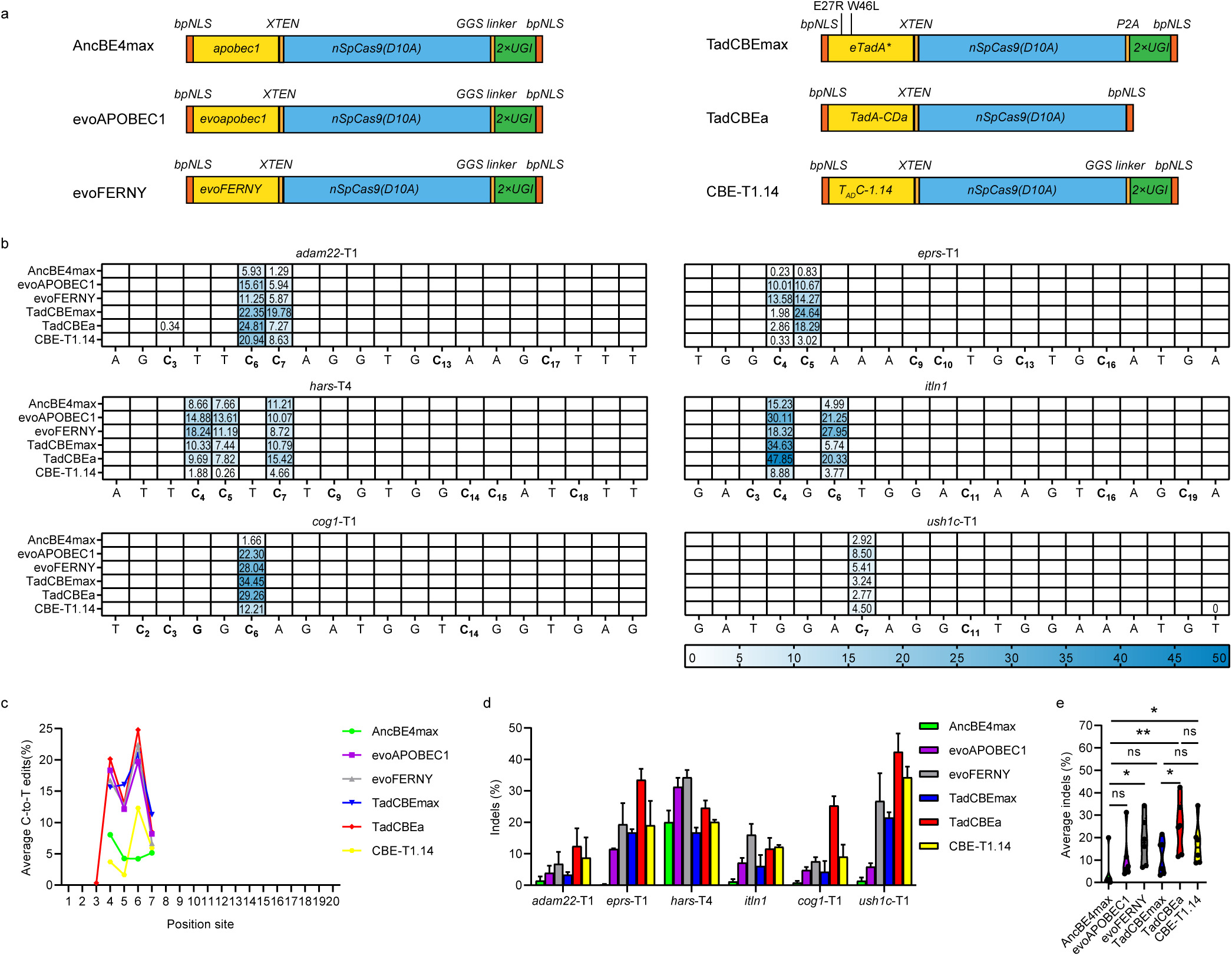
Comparative evaluation of in vivo editing efficiencies for six cytosine base editors in zebrafish. **(a).** Schematics of the constructs for six cytosine base editors. bpNLS: bipartite nuclear localization, Apobec1, evoApobec1, evoFERNY, eTadA*, TadA-CDa and TADC-1.14: various adenine deaminase, XTEN: a 32aa flexible linker, nSpCas9: SpCas9 nickase, GGS linker: GGSSGGS amino acid, P2A: Porcine teschovirus-1 2A, UGI: Uracil glycosylase inhibitor. **(b).** The C-to-T editing efficiency of AncBE4max, evoAPOBEC1, evoFERNY, TadCBEmax, TadCBEa, CBE-1.14 was examined at 6 endogenous genomic loci. The heatmap represents the average editing percentage derived from three independent experiments. The color mapping from blue to white represents the editing efficiency from 50% to 0%. **(c).** Frequencies of C to T editing across protospacer at the 6 sites (PAM located at positions 21–23). Single dots represent the average C-to-T editing efficiency at each position. **(d).** The indel efficiency comparison among six cytosine base editors targeting six different loci. Values are presented as mean value ± standard deviation (SD), n = 3 biological replicates. Data are expressed as mean ± SD. **(e).** Assessment of the mean indel efficiency of six cytosine base editors using the plot based on the data in Fig. 1d. Mean indel frequency at each site is represented by individual data points, with the central dotted line indicating the overall mean. Two-tailed paired t-test were performed (with P values marked at the top of the violin plot.)

### Characterization of a New TadA-Derived CBE with High Efficiency and Low Indel Frequency

Given TadCBEa’s high on-target activity and TadCBEmax’s low indel frequency, we hypothesized that a new cytosine base editor (CBE) system could be constructed by integrating their advantageous features. Notably, TadCBEa lacks a uracil glycosylase inhibitor (UGI) motif, while TadCBEmax consists of one. In the context of cytosine base editing, the UGI motif is essential for inhibiting uracil DNA glycosylase (UDG), which typically reverts deaminated cytosine (uracil) to cytosine as part of a DNA repair mechanism. By blocking UDG, UGI therefore enables the stable conversion of uracil to thymine, thereby enhancing the efficiency, accuracy, and stability of the intended C-to-T edit^1^. Recently, we developed an advanced TadA-based adenine base editor (ABE) variant, ABE-ultramax, which includes V82S and Q154Q mutations in the deaminase domain. This variant achieved up to 100% efficiency in A-to-G editing in zebrafish^23^. We hypothesized that these beneficial mutations might similarly enhance TadA-derived CBE systems. Accordingly, we introduced the V82S and Q154Q mutations, together with two copies of UGI connected by a Glycine-Glycine-Serine (GGS) linker, to the C-terminal of CRISPR-associated protein 9 (Cas9), creating a new CBE variant that we named zTadCBE (**Fig. 2a**). Comparative analyses across the six target loci revealed that zTadCBE exhibited 1.3-fold and 2-fold higher on-target editing efficiency compared to TadCBEa and TadCBEmax, respectively (**Fig. 2b and 2c**). Further, indel frequency in zTadCBE was comparable to the low indel rates of TadCBEmax and were significantly reduced relative to TadCBEa (**Fig. 2d and 2e**).

**Fig. 2:**
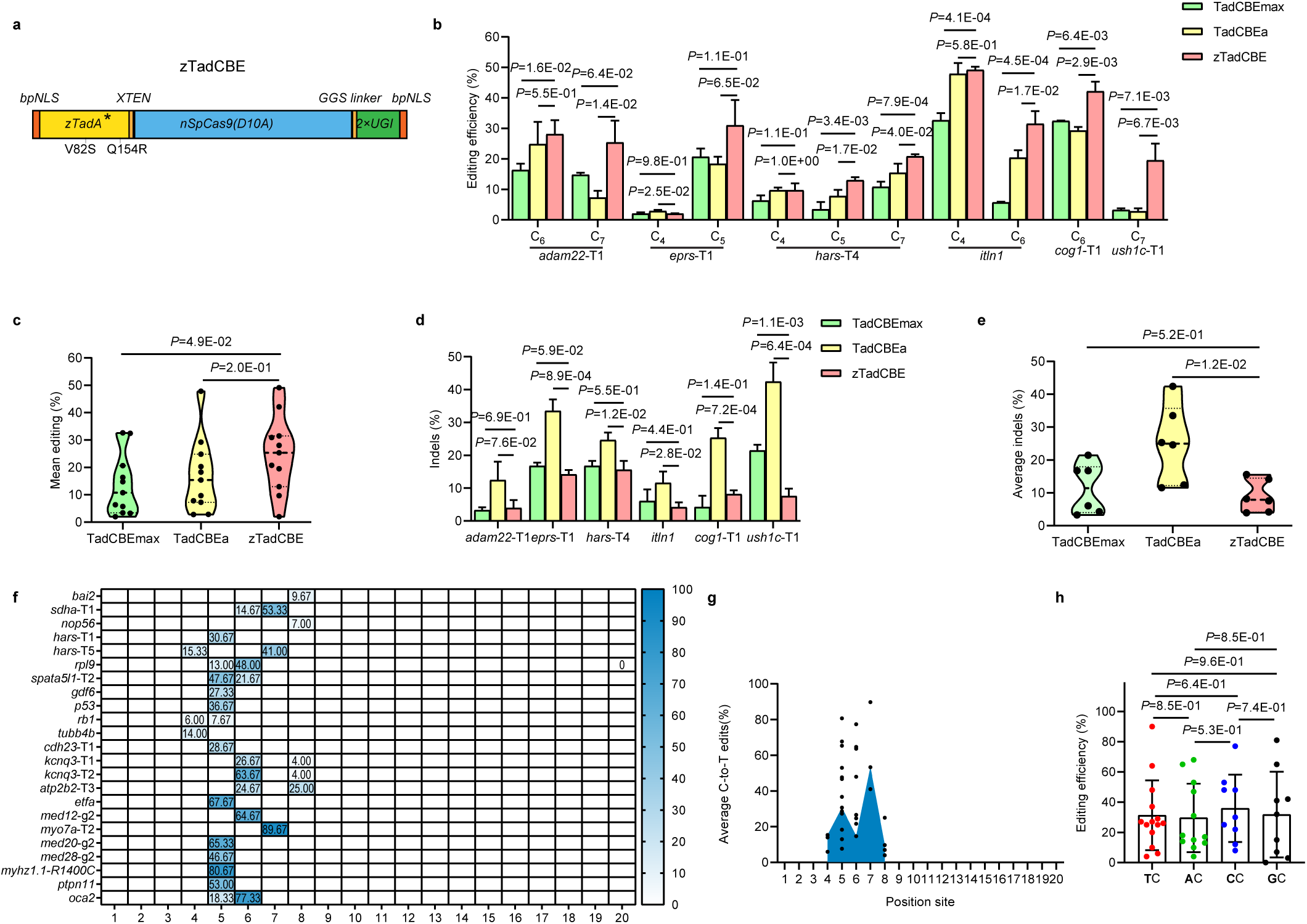
Efficient cytosine base editing mediated by zTadCBE in zebrafish. **(a).** Schematics of the zTadCBE construct, featuring the new zTadA* variant that incorporates TadCBEa mutations along with two additional mutations, V82S and Q154Q. bpNLS: bipartite nuclear localization, XTEN: a 32aa flexible linker, nSpCas9 (D10A): SpCas9 nickase, GGS linker: GGSSGGS amino acid, UGI: Uracil glycosylase inhibitor. The prefix “z” denotes zebrafish-specific codon optimization. **(b).** Comparison of editing efficiency among TadCBEmax, TadCBEa, and zTadCBE across six target loci. Base position within the sgRNA is denoted numerically, and values are reported as mean ± standard deviation (SD), with n = 3 biological replicates. Statistical analysis was conducted using a two-tailed paired t-test, with P values indicated. **(c).** Analysis of mean editing efficiency for TadCBEmax, TadCBEa, and zTadCBE based on data in Fig. 2b. Mean editing efficiency per site is shown by individual data points, with the central dotted line representing the overall mean. Two-tailed paired t-tests were performed, with P values annotated at the top of the violin plot. **(d).** Comparison of indels efficiencies among TadCBEmax, TadCBEa, and zTadCBE across six target loci. Values are reported as mean ± standard deviation (SD), with n = 3 biological replicates. Statistical analysis was conducted using a two-tailed paired t-test, with P values indicated. **(e).** Analysis of mean indels frequency for TadCBEmax, TadCBEa, and zTadCBE based on data in Fig. 2b. Mean editing efficiency per site is shown by individual data points, with the central dotted line representing the overall mean. Two-tailed paired t-tests were performed, with P values annotated at the top of the violin plot. **(f).** Heatmap illustrating the average C-to-T editing efficiency of zTadCBE across 23 target sites. Editing efficiency is shown on a color scale, where blue represents 100% efficiency and white represents 0% efficiency. **(g).** Evaluation of zTadCBE efficiency and targeting window. Each data point reflects the mean editing activity at a specific site. The targeting window of zTadCBE, indicated in blue, spans from position 4 to 8, counted from the 5’ to 3’ ends of the target site. Data from three independent experiments were analyzed. **(h).** Base editing efficiencies of the zTadCBE systems at the target C in different sequence contexts. Each data point reflects the mean editing activity at a specific site. Statistical analysis was conducted using a two-tailed paired t-test, with P values indicated.

To comprehensively assess the performance of zTadCBE in zebrafish, we examined 23 additional endogenous targets. Editing activity, defined as the maximum observed activity at any position within each target, ranged from 7.67% to 89.67% (**Fig. 2f**). The primary editing window of zTadCBE spans five nucleotides (positions 4–8) (**Fig. 2g**), consistent with the editing window of AncBE4max^8^. We further observed that zTadCBE did not exhibit a pronounced sequence context bias (**Fig. 2h**), although reduced activity was noted at specific AC or GC sites.

To assess Cas9-dependent off-target activity of zTadCBE in zebrafish, we analyzed the top three off-target sites predicted for three loci by the CRISPOR tool. We PCR-amplified each target and subjected them to next-generation sequencing. Our data indicated that off-target editing at these sites was minimal, remaining nearly undetectable (<3%) (**Table 1**). Previous studies have reported that TadA-derived CBEs also possess some A-to-G editing activity, leading us to further evaluate 20 loci for A-to-G conversions^18–20^. Our results indicated that 4 out of these 20 sites exhibited minimal A-to-G activity (<10%) (**Supplementary** Fig. 1), with most edited A bases situated within TA motifs, aligning with TadA’s intrinsic TA-site preference. Collectively, our findings suggested that zTadCBE, an engineered cytosine deaminase variant, is a promising tool for achieving high efficiency and precision in C-to-T base editing in zebrafish, with minimal off-target effects.

**Table1.** Assessment of on-target and off-target editing by zTadCBE using NGS in zebrafish. On-target, product purity and off-target analysis of zTadCBE induced C-to-T editing at *p53*, *kcnq3*-T2 and *spata5l1*-T2 sites using NGS. The top three high-scoring off-target sites (PAM are underlined) are shown. Mismatched bases are indicated in lowercase. The PAM sequences are underlined in red

### Expanding the Targeting Range and Editing Windows of TadA-derived CBEs

Base editors derived from the CRISPR-associated protein 9 of *Streptococcus pyogenes* (SpCas9) are limited by the NGG protospacer-adjacent motif (PAM), restricting base editing to an editing window usually 4-8 bases distal to the PAM. Recent studies have demonstrated that the engineered SpCas9 variant, SpRY, with highly flexible PAM sequences, is compatible with single-base editing systems in zebrafish, enabling the recognition of atypical NNN PAM sequences and thus expanding the targetable range of base editors^3,24^. To broaden the targeting scope of zTadCBE-mediated cytosine base editing, we substituted SpCas9n of the zTadCBE construct with SpRYnCas9, resulting in zTadCBE-SpRY. To assess the C-to-T editing efficiency of zTadCBE-SpRY at NNN PAM sites in zebrafish, we selected 12 target loci with atypical PAM sequences and injected single guide RNAs (sgRNAs) together with zTadCBE-SpRY-encoding mRNA into one-cell stage embryos. Sequencing results showed that all 12 loci exhibited C-to-T conversion with variable efficiencies across all sites (**Fig. 3a**). In comparison to CBE4max-SpRY—the most flexible cytosine base editor (CBE) reported thus far for targeting the zebrafish genome^25^—zTadCBE-SpRY demonstrated an average six-fold increase in C-to-T editing efficiency (**Fig. 3a and 3b**). Importantly, it could also effectively target specific loci inaccessible to CBE4max-SpRY, such as rpl18-NAC, rps16-NAA, and med20-NAA (**Fig. 3a**). These findings indicated that TadA-based C-to-T base editing was compatible with SpRYCas9n, supporting efficient editing by recognizing a highly versatile PAM range in zebrafish, further expanding the targeting range.

**Fig. 3:**
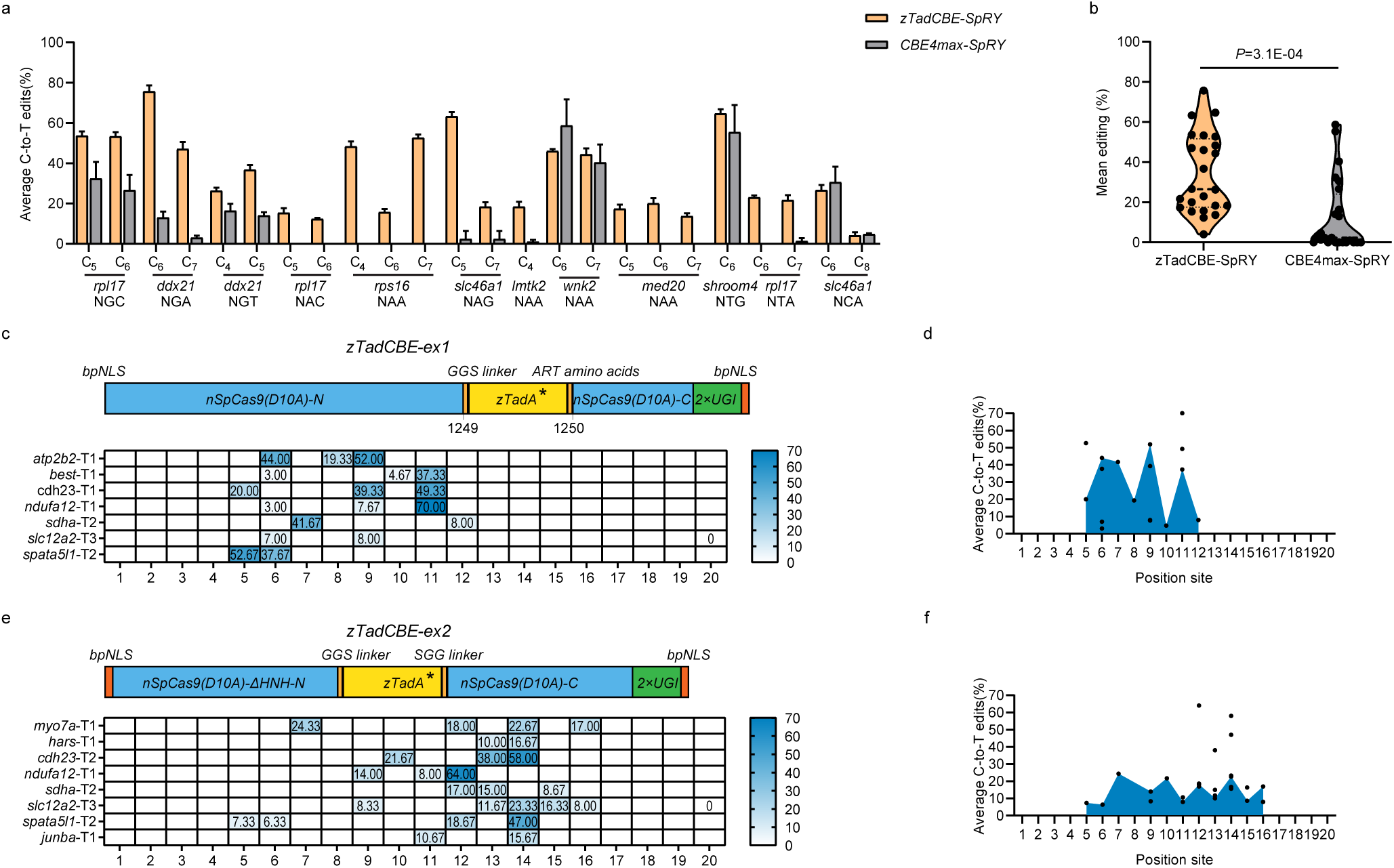
Expanding Targeting Range and Editing Windows by zTadCBE variants. **(a)**. Comparison of editing efficiencies between zTadCBE-SpRY and CBE4max-SpRY Using twelve gRNAs Targeting NNN PAMs. The position of the edited base within each sgRNA is indicated numerically. Data are presented as mean values ± standard deviation (SD), calculated from three biological replicates. **(b)**. Analysis of mean editing efficiency for zTadCBE-SpRY and CBE4max-SpRY based on data in Fig. 3a. Mean editing efficiency per site is shown by individual data points, with the central dotted line representing the overall mean. Two-tailed paired t-tests were performed, with P values annotated at the top of the violin plot. **(c)**. Schematics showing constructs of zTadCBE-ex1 designed to shift the editing window of cytosine base editing.Heatmap illustrating the average C-to-T editing efficiency of zTadCBE-ex1 across seven target sites. Editing efficiency is shown on a color scale, where blue represents 70% efficiency and white means 0% efficiency. **(d)**. Evaluation of zTadCBE-ex1 efficiency and targeting window. Each data point reflects the mean editing activity at a specific site. The targeting window of zTadCBE-ex1, indicated in blue, spans from position 5 to 12, distal from the PAM. Data from three independent experiments were analyzed. **(e)**. Schematic Representations of zTadCBE-ex2 Constructs. A heatmap depicting the average C-to-T editing efficiency of zTadCBE-ex2 across eight target sites is shown, with a color gradient indicating efficiency levels, where blue denotes 70% efficiency and white indicates 0% efficiency. **(f)**. Assessment of zTadCBE-ex2 efficiency and targeting window. Each data point corresponds to the mean editing activity observed at a specific target site. The targeting window for zTadCBE-ex2, highlighted in blue, extends from position 5 to 16, distal from the PAM. Data were analyzed from three independent experiments.

While the PAMless variant could also expand the targeting range, the editing window of zTadCBE-SpRY or zTadCBE remains limited. This constrained editing window is a significant limitation to the broad applicability of single-base editing tools. We and others have previously shown that altering the relative positioning between the deaminase and Cas9 can shift the target window of adenine base editors (ABEs)^23,26,27^. Whether this approach could also be applied effectively to TadA-based CBEs has not yet been reported. To explore this possibility, we used the same positioning strategy to construct two novel CBE variants, zTadCBE-ex1 and zTadCBE-Ex2 (**Fig. 3c and 3e**), and tested their target windows using seven or eight target sites in zebrafish, respectively. zTadCBE-ex1 primarily edited cytosine bases between positions 5 to 12 of the protospacer (distal to the PAM) (**Fig. 3c and 3d**). In contrast, zTadCBE-ex2 shifted the editing window closer to the PAM-proximal end of the protospacer—positions 5 to 16 (**Fig. 3e and 3f**)—as compared to positions 4–8 typically targeted by conventional CBEs.

Taken together, our findings demonstrated that the PAM-flexible variant, zTadCBE-SpRY, together with the zTadCBE-ex1 and zTadCBE-ex2 variants, provide high-efficiency, expanded C-to-T editing. This alleviates the previously restrictive editing capabilities of CBEs and provides access to previously un-targetable sites, offering greater flexibility in selecting efficient editors for diverse loci of interest. Our observation of consistently high (≈ 60%) germline targeting efficiencies, with transmission rates of >70% for both zTadCBE and zTadCBE-SpRY across four distinct loci (**Supplementary Table 1**), further confirmed their ability to transmit targeted edits to the germline robust capacity to produce precise base edits with substantial efficiency.

### Disease Modeling in Zebrafish Using TadA-Derived CBEs

Next, we aimed to utilize zTadCBEs to generate disease-related genetic variants of zebrafish, leveraging the unprecedented ability of cytosine base editors (CBEs) to introduce patient-specific point mutations that mimic the genetic basis of many human diseases^28^.

First, we focused on hearing loss—a prevalent congenital sensory impairment, a significant proportion of cases of which are identified prior to speech development (prelingual). This condition encompasses a substantial level of genetic heterogeneity— genetic factors account for approximately 80% of prelingual hearing loss cases, which frequently also lack any physical symptoms (non-syndromic hearing loss)—an auditory phenotype associated with over 150 genes^29^. Among these genes, *CDH23* stands out as a prominent contributor. This gene encodes cadherin 23, a cell adhesion protein crucial for the proper development and function of the inner ear and retina. Given its critical role, mutations in *CDH23* can lead to a spectrum of auditory phenotypes, from nonsyndromic autosomal recessive deafness-12 (DFNB12) to complex genetic disorders characterized by combined deafness and blindness as seen in the severe Usher syndrome type 1D (USH1D)^30^.

To study the effects of patient-specific genetic mutation in *CDH23,* we developed a disease model targeting the c.6085C>T (p.Arg2029Trp) variant of the *CDH23* gene, reported in patients with non-syndromic hearing loss^31^. To model this variant, we relied on the zebrafish inner ear and specialized organ system, the lateral line system, which share structural and functional similarities with the human auditory system, making them effective models for studying hearing and balance. In particular, the zebrafish lateral line system in accessible and contains functionality similar hair cells to the inner ear hair cells, and transparent embryos provided us with a valuable tool for studying the function and development of hair cells—the primary sensory receptors of the inner ear—using live dyes such as FM1-43, DASPEI, and Yo-Pro-1^32^. To target the base to test this variant, we selected a single guide RNA (sgRNA) site with an NGT protospacer-adjacent motif (PAM) (**Fig. 4a**). The target cytosine was located within a CC sequence context, which could not be edited by APOBEC1-based deaminases due to sequence context preferences. Using zTadCBE-SpRY, we could target this cytosine (position 7 from the 5′ end) with a mean editing efficiency of 48.27% ± 2.46%. This also resulted in editing a bystander cytosine at position 6 (**Supplementary Fig. 2a**), leading to a synonymous mutation that did not alter the amino acid sequence of *cdh23*. Given the TC sequence context preference of APOBEC-based base editors, as expected, CBE4max-SpRY was unable to edit this site (**Supplementary Fig. 2a**). After raising the edited animals to adults, we identified mutations in the F1 generation with 100% efficiency, and further analyzed phenotypes in the F2 generation by breeding two F1 carriers. We observed that homozygous larvae failed to take up the live dye Yo-Pro-1, which is typically taken up by the mechanotransduction channels of hair cells. Our results in zebrafish models confirmed the utility of TadA-Derived CBEs for studying the functional consequences of clinically relevant human single nucleotide variants (SNVs), such as those of the *CDH23* gene demonstrated here (**Fig. 4b and 4c**).

**Fig. 4:**
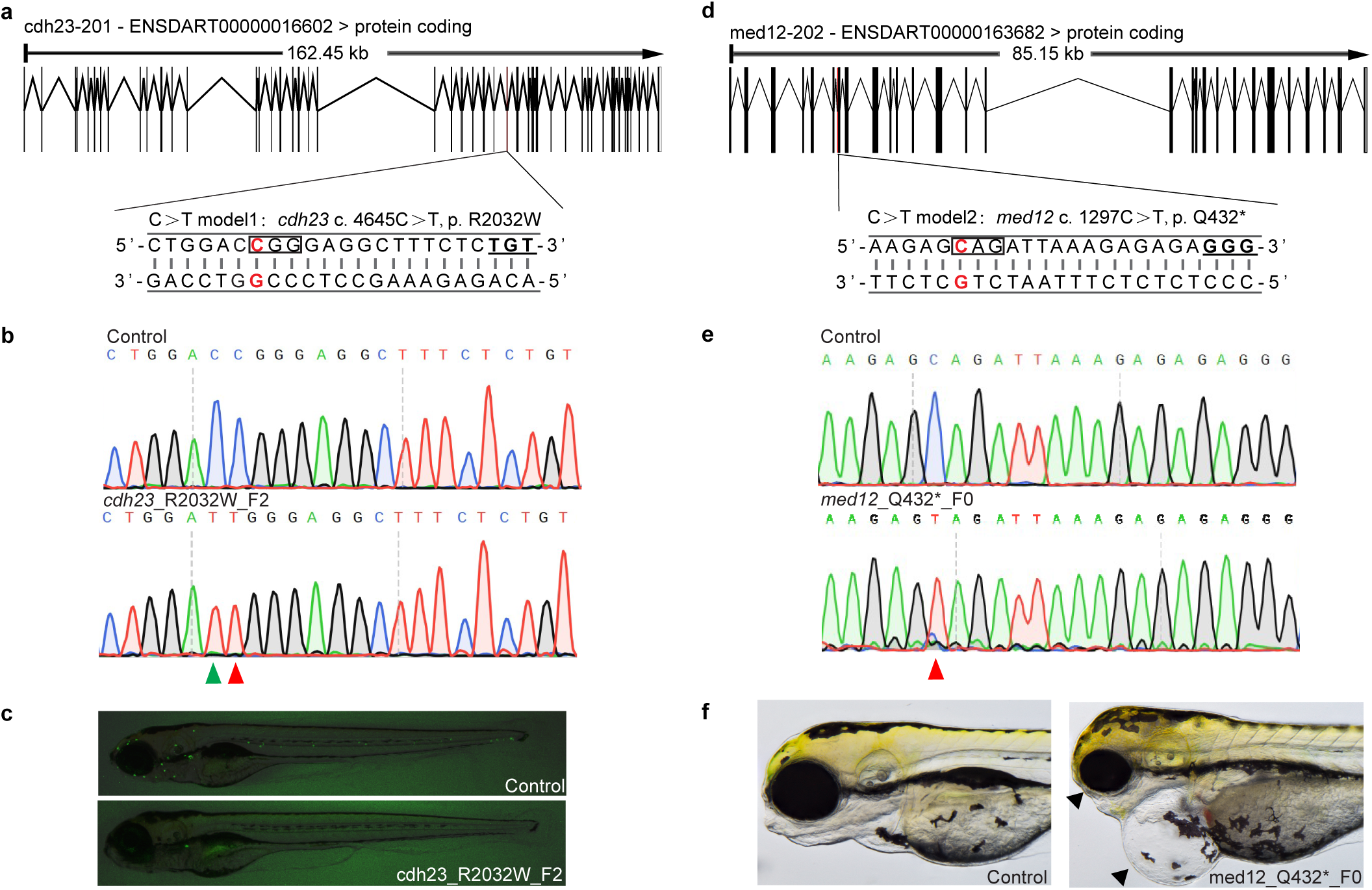
Functional assessment of genetic variants using zTadCBE editors. **(a)**. Schematic diagrams of *cdh23* (R2032W) in zebrafish. The target sequence is displayed with the PAM underlined and bold. The targeted nucleotide is highlighted in red, the related coding frame is indicated by a black box. **(b)**. Sequencing results of *cdh23* (R2032W) F2 embryos induced by zTadCBE-SpRY. The red arrowhead points to the expected nucleotide substitutions, while a green arrowhead in the Sanger sequencing chromatograms indicates bystander base substitutions. **(c)**. Phenotype of *cdh23* (R2032W) F2 embryos with mutation induced by zTadCBE-SpRY. Compared to the control, F2 *cdh23* (R2032W) embryos exhibited a significant reduction in functional neuromast hair cells (labeled with Yo-Pro1 (green)). **(d)**. Schematic diagrams of *med12* (Q4322*) in zebrafish. The targeted sequence is displayed with the PAM underlined and bold. The targeted nucleotide is highlighted in red, the related coding frame is indicated by a black box. **(e)**. Sequencing results of *med12* (Q4322*) F0 embryos induced by zTadCBE. The red arrowhead points to the expected nucleotide substitutions. **(f)**. Phenotypes of *med12* (Q4322*) F0 embryos induced by zTadCBE. Compared to the control, *med12* (Q4322*) F0 embryos exhibited microcephaly, microphthalmia, and pericardial edema (black arrowhead) phenotypes.

We further induced point mutation in the *MED12* gene, encoding Subunit 12 of the Mediator complex that regulates transcription by facilitating interactions between transcription factors and RNA polymerase II. Given its essential role, mutations in *MED12* are implicated in various developmental and neuropsychiatric disorders^33^. CRISPR-based selective targeting of optimal polymorphisms (CRISPR-STOP) has recently proven to be an efficient, less disruptive alternative to traditional Cas9 approaches in gene-knockout applications^34^. Using this strategy, we designed a sgRNA to induce a premature termination codon, generating a *med12* mutant in zebrafish. The selected target site for this mutation was located within a GC sequence context (**Fig. 4d**), which the zTadCBE could specifically edit with nearly 100% editing efficiency (**Fig. 4e**). In contrast, the AncBE4max CBE failed to produce any edits at this site (**Supplementary Fig. 2b**), underscoring the efficiency and adaptability of TadA-derived zTadCBE in GC-rich contexts. Further phenotypic analysis of F0 embryos injected with zTadCBE and the corresponding sgRNA revealed developmental abnormalities that were consistent with previously reported *med12* mutant models, including microcephaly, microphthalmia, and pericardial edema (**Fig. 4f**)^35^. Collectively, our findings suggested that highly efficient CBEs are promising tools for rapid F0-based functional gene analysis. Compared to previous tools, zTadCBE and related variants demonstrated substantial advantages in both editing efficiency and target selectivity, making them particularly well-suited for constructing zebrafish disease models, greatly enhancing the precision and flexibility of genetic research in this area.

## Discussion

Given the rapidity at which large numbers of genetic mutations are being identified, the development of clustered regularly interspaced short palindromic repeats (CRISPR) base editors was a significant advancement in CRISPR technology, allowing researchers to modify single nucleotides within the genome—particularly disease-specific mutations— with high precision and efficiency. As such, base editors hold immense promise for studying disease-specific variants in genetically tractable, mammalian-like model organisms such as mice and zebrafish.

Cytosine base editors (CBEs) based on APOBEC1 deaminases such as BE3 and BE4^1^ laid the groundwork for the continued development of base editors. However, their utility in animal models has been historically constrained due to limitations in considerable off-target effects and sequence biases—particularly in zebrafish where TC-context biases are more acute than in mammalian cells^36^—limiting their editing capabilities in CC and GC contexts. While recent advancements, like AncBE4max, have improved the editing efficiency and accuracy of base editors^8^, they continue to exhibit sequence-specific constraints and produce more insertions and deletions (indels) in zebrafish.

In this study, we developed an optimized TadA-derived CBE for zebrafish, zTadCBE, which addresses these limitations by significantly improving editing efficiency and sequence bias, minimizing off-target edits, and expanding target site accessibility. We further demonstrated zTadCBE’s potential to greatly broaden the application of CBEs to target specific sequence contexts, by applying it to successfully construct zebrafish disease models with edited CC and GC loci. This paves the way for functional screening of genetic variants *in vivo* in zebrafish.

One major challenge in applying single-base editors to zebrafish is maintaining high precision. Existing methods, including linker modifications between the base editor’s structural components—the deaminase and the CRISPR-associated protein 9 (Cas9)— or targeted mutations within the deaminase, have been reported to enhance base editor precision^37,38^. However, whether these approaches can be adapted to improve zTadCBE remains to be studied. Our finding of minimal off-target edits for zTadCBE align with previously reported low off-target effects of TadA-based adenine base editors (ABEs)^23^, showing that TadA-derived CBEs also exhibit high specificity in zebrafish, which could hold substantial clinical value. Despite these advantages, zTadCBE’s efficiency still varies across different loci.

Further enhancements to zTadCBE to achieve higher editing activity in the F0 generation could significantly reduce the time required for assessing single nucleotide variant (SNV) pathogenicity, like what we have achieved with our ABE-Umax variant^23^. Recently, David Liu’s lab identified next-generation CBE6 variants via phage-assisted evolution^39^, and Erwei Zuo’s group developed the CD0208 base editor using structure-based generative machine learning, demonstrating superior performance in cellular models^40^. Future work will investigate whether these advanced tools can be adapted in zebrafish, potentially enabling biallelic editing in the F0 generation for accelerated functional studies.

## Methods

### Ethical Statement

All zebrafish experiments were carried out as per protocols 24-11, 24-12 and 24-28 approved by the Institutional Animal Care Committee (IACUC) of OMRF.

### Zebrafish maintenance

Wildtype zebrafish strain NHGRI-1^41^ were raised and maintained at 28.5 °C on a 14 h light/10 h dark cycle. The selection of mating pairs (12-15 months) was random from a pool of 30 males and 30 females. All experiments were conducted using wild-type zebrafish.

### Construction of Plasmids

All variant plasmids were constructed based on the pT3TS-zevoCDA1-BE4max plasmid^37^ with respective modifications. For the AncBE4max, evoAPOBEC1, evoFERNY, TadCBEmax, TadCBEa, and CBE-T1.14 plasmids, deaminase sequences were optimized to follow zebrafish codon preferences and were synthesized by GenScript. The optimized deaminase sequences replaced the original zevoCDA1 sequence. In the TadCBEa plasmid, the downstream 2X UGI sequence was removed. For the TadCBEmax construct, the GGS linker was substituted with a P2A sequence. For the zTadCBE plasmid, we introduced two mutations, V82S and Q154R, in the deaminase region of TadCBEa and added 2X UGI at the C-terminus of Cas9. For zTadCBE-SpRY, the SpRY sequence was cloned from the ABE-Umax-SpRY plasmid and replaced the SpCas9 component in zTadCBE. In constructing the zTadCBE-ex1 plasmid, the zTadA* variant was inserted at the SpCas9 docking site at position 1249, using a GSSGSS linker and an ART amino acid linker to connect the N- and C-termini of zTadA* with Cas9. For zTadCBE-ex2, the HNH domain of Cas9 was removed, and a GGS linker was used to link SpCas9 S793 to the N-terminus of zTadA*, with an SGG linker connecting the zTadA* C-terminus to SpCas9 R919. All fusions and mutations were generated using the Vazyme Mut Express II Fast Mutagenesis Kit V2 (Cellagen Technology LLC, CA, USA). The assembled plasmids were then transformed and amplified in DH5α Chemically Competent Cells (New England Biolabs, MA, USA).

### Single guide RNAs (sgRNAs) and mRNA synthesis

All sgRNAs used in this study were chemically modified with 2′-O-methyl (M) and 2′-O-methyl 3′-phosphorothioate (MS) modifications at both the 5’ and 3’ ends. They were synthesized by GenScript Inc. (NJ, USA) and Synthego Inc. (CA, USA). Detailed target sequences are provided in **Supplementary Table 2.** All mRNAs were transcribed in vitro from XbaI-linearized templates (NEB, USA) using the T3 mMESSAGE mMACHINE Kit (ThermoFisher Inc., CA, USA) and purified with the Monarch RNA Cleanup Kit (New England Biolabs, MA, USA). The resulting capped mRNAs and synthetic sgRNAs were prepared in a 2000 ng/μl stock solution and stored at -80°C for subsequent use.

### Embryo microinjection, morphological phenotyping, and imaging

A solution containing sgRNA at 200 ng/μl and Cas9 mRNA at 400 ng/μl was injected in a 2-nl volume into one-cell-stage embryos. After 2 to 5 days post-fertilization (dpf), the embryos were anesthetized in a 0.016% tricaine/MS-222 solution (Sigma-Aldrich, MO, USA) and positioned in 3% methylcellulose (Sigma-Aldrich, MO, USA) for imaging. Neuromast hair cells were visualized at 5 dpf by incubating the embryos in 1 mM YO-PRO™-1 (Invitrogen, USA) dye for one hour at room temperature. Images were captured using an Olympus SZX12 stereomicroscope equipped with a DP71 color digital camera (Olympus, Japan). Following imaging, the embryos were genotyped to establish connections between their genotypes and observable phenotypes.

### Base editing analysis

Genomic DNA was extracted from three pooled samples, each containing six randomly selected embryos. A targeted locus of 150-300 bp was amplified using HotStart Taq-Plus DNA polymerase (Qiagen, USA) with primers listed in **Supplementary Table 2**. The PCR products were purified using the DNA Clean & Concentrator-5 kit (ZYMO, USA) for sequencing. For Sanger sequencing, the results were analyzed with EditR (v1.0.10)^42^ (31021262). For Next-Generation Sequencing (NGS), paired-end sequencing was performed on the Illumina MiSeq platform, and the data were processed using CRISPResso2 software to assess editing outcomes^43^.

### DNA Off-target analysis

To identify potential off-target effects, each guide RNA (gRNA) underwent an in silico off-target prediction analysis using Cas-OFFinder (24463181) and CRISPOR (Version 4.99)^44^. The specificity score for each predicted off-target site was calculated with CRISPOR. The top three off-target sites with the highest specificity scores were then selected for further validation using next-generation sequencing (NGS).

### Statistics & Reproducibility

Sample sizes were not predetermined through statistical methods; however, randomization was applied, and no data were excluded from analysis. Each experiment was repeated at least three times, with sample sizes detailed in the figure legends or Source Data. Data are shown as the mean ± standard deviation (SD). Statistical analyses were performed using GraphPad Prism version 8.0.2 (GraphPad Software, USA). Statistical significance was defined as *p < 0.05, **p < 0.01, and ***p < 0.001. To compare base editing efficiencies between different groups, two-tailed unpaired Student’s t-tests were conducted, while comparisons of mean editing efficiencies within groups were evaluated using two-tailed paired Student’s t-tests or nonparametric Wilcoxon matched-pairs signed rank tests.

## Data Availability

NGS data are available on the National Center for Biotechnology Information Sequencing Read Archive (SRA) database under project number (https://www.ncbi.nlm.nih.gov/bioproject/PRJNA1118438) and. All data supporting the findings of this study are available within the article and Supplementary Information files. Source data are provided in this paper. Plasmids from this study are available at Addgene.

## Acknowledgments

This work was supported by US National Institutes of Health (NIH) grant 1R24OD034438 (G.K.V.) and Presbyterian Health Foundation (G.K.V.).

## Author contributions Statement

W.Q., and G.K.V. conceived the project. W.Q. performed the experimental work and analyzed the results. S.L., C.P., and P.V. contributed to the experiments. W.Q, and G.K.V. wrote the original draft. All other authors reviewed and edited the manuscript. G.K.V. acquired funding and supervised the study.

---

**Supplementary Figure 2.** Sanger sequencing results of *cdh23* (R2032W) and *med12* (Q4322*) F0 embryos induced by different cytosine base editors.

**(a)** Sanger sequencing results of *cdh23* (R2032W) F0 embryos induced by CBE4max-SpRY and zTadCBE-SpRY, respectively.

**(b)** Sanger sequencing results of *med12* (Q4322*) F0 embryos induced by ancBE4max and zTadCBE, respectively.

The red arrowhead points to the expected nucleotide substitutions, while a green arrowhead in the Sanger sequencing chromatograms indicates bystander base substitutions.

